# Integrating mass spectrometry with Nanopore direct RNA sequencing for *de novo* modification profiling of bacteriophage MS2

**DOI:** 10.64898/2026.06.01.729303

**Authors:** Joshua D Jones, Neda Ghohabi Esfahani, Calvin LC Goemann, Tanner Wiegand, Artem Nemudryi, Minli Ruan, Xiaoyan Li, Mehmet Tardu, Robert T Kennedy, Blake Wiedenheft, Miten Jain, Kristin S Koutmou

## Abstract

RNA modifications contribute to the regulatory and structural complexity of RNA molecules, influencing key biological processes. Nanopore direct RNA sequencing (DRS) enables the detection of these modifications at single-molecule resolution without chemical conversion. Oxford Nanopore Technologies recently introduced RNA004 sequencing chemistry and the Dorado basecaller, which improves accuracy and enables the identification of eight RNA modifications. However, the reliability of these predictions requires careful validation using orthogonal approaches. Here, we profiled RNA modifications in the MS2 bacteriophage genome using both LC-MS/MS and Nanopore DRS. LC-MS/MS analysis revealed that Ψ is the predominant modification, present at approximately one site per RNA molecule, while all other modifications were found at low levels (< 0.04 modifications per RNA). In contrast, Dorado-based modification calling not only confirmed Ψ but also predicted multiple m^6^A, m^5^C, 2’OMe, and inosine sites not supported by LC-MS/MS data. To refine modification calling, we used in vitro transcribed (IVT) RNA as a negative control and subtracted IVT-derived false positive rates from native RNA predictions. This adjustment reduced overcalling and improved confidence in site-specific predictions. These data demonstrate that while Dorado can detect RNA modifications *de novo*, its predictions require careful filtering and validation. Our studies of provide a rare benchmark dataset for assessing the ability of transcriptome-wide methods to accurately identify RNA modifications and estimate modification stoichiometry. These findings support the use of orthogonal approaches, including LC-MS/MS and IVT controls, alongside Nanopore sequencing to provide a more reliable and interpretable strategy for studying RNA modification patterns.

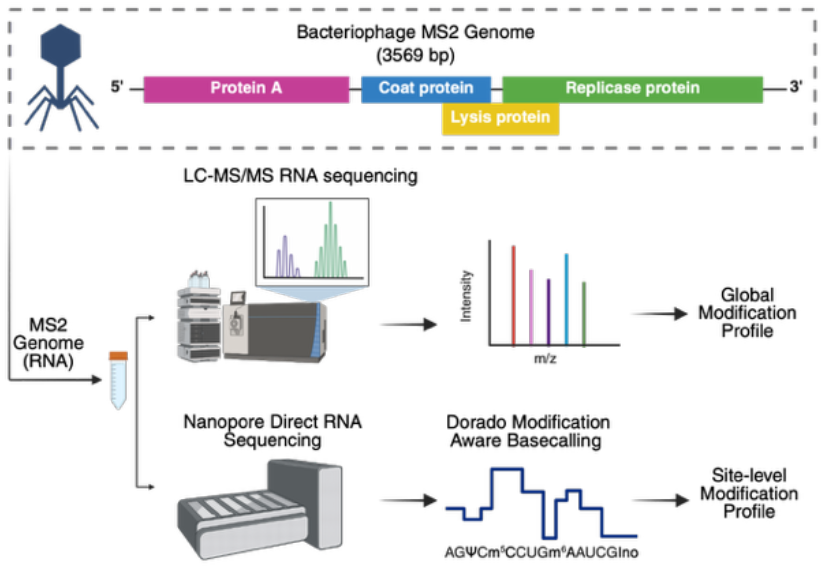

## INTRODUCTION

RNA modifications are widespread chemical alterations that expand the functional complexity of the transcriptome far beyond adenosine, guanosine, cytidine and uridine. There are > 170 known varieties of modifications that make key contributions to RNA stability, structure, and translation ^1–3^. Modifications are especially abundant in three core components of the protein synthesis machinery: ribosomal RNA (rRNA), transfer RNA (tRNA) and messenger RNA (mRNA) ^4^. As such, it is unsurprising that RNA modifications are implicated in modulating gene expression and cellular response mechanisms ^1,5,6^. Despite their importance, the exact amounts and precise locations of most RNA modifications are not known across biology. This is largely due to long-standing technical challenges in the detection and mapping of RNA modifications. These difficulties arise from their relatively low abundance - often 20-to 10,000-fold lower than canonical nucleosides - and sub-stoichiometric incorporation ^7,8^.

RNA modification detection methods generally fall into two complementary categories: nucleoside detection and modification mapping. Global ribonucleoside modification profiling methods largely rely on liquid chromatography tandem mass spectrometry (LC–MS/MS) to provide sensitive, multiplexed and quantitative measurements of total modification content within a purified RNA sample ^7,9–11^. Because RNA is enzymatically digested into nucleosides before analysis, nucleoside LC-MS/MS approaches provide information on modification identity and abundance, but not on transcriptome-wide location ^7,11^. Therefore, orthogonal methods are required to map modifications. LC–MS/MS-based sequencing is the gold-standard strategy for RNA modification mapping, but it requires RNA inputs that are prohibitive for most organisms and experimental systems, exceeding 10 μg ^12–14^. Next-generation RNA sequencing methods are beginning to address this limitation, though they often rely on an intermediate chemistry step (e.g., reverse transcription) prone to bias, and typically detect a limited chemical variety of modifications within a single experiment ^15–19^. In contrast, Nanopore direct RNA sequencing (DRS) sequences native RNA molecules directly and can, in principle, detect nucleotide modifications without chemical conversion, reverse transcription or amplification ^20,21^.

Nanopore DRS is quickly emerging as a platform that can potentially provide transcriptome-wide, site-specific information about the position of numerous modifications. DRS measures changes in ionic current as RNA molecules translocate through a biological nanopore. These ionic current signals are decoded into nucleotide sequences using deep-learning models. Over the past decade, improvements in Nanopore DRS have enabled reliable sequencing of canonical ribonucleotides, detection of altered modification status at known sites in cellular RNAs ^22–25^, identification of RNA-modifying enzymes associated with specific sites ^26,27^ and characterization of RNA isoforms in cells ^28^. Nanopore DRS also requires relatively low (∼1 μg) total RNA input ^29^, making it a powerful and rapidly emerging central platform for RNA modification analysis.

Until recently, most DRS studies used ONT’s RNA002 platform. In 2024, ONT released the upgraded RNA004 platform together with the Dorado basecaller, which improved basecalling accuracy and expanded modification-detection capabilities ^30^. Dorado now supports *de novo* calling of eight RNA modifications, including pseudouridine (Ψ), N6-methyladenosine (m^6^A), 5-methylcytidine (m^5^C) and inosine. However, the performance of these modification calls has not yet been systematically evaluated against orthogonal biochemical methods. Several semi-quantitative Nanopore-based approaches have been developed using synthetic oligonucleotides, but these controlled substrates do not fully capture the complexity of biological RNA ^22,31–35^. Unlike conventional RNA-seq, Nanopore ionic-current signals reflect short sequence contexts, typically k-mers of approximately five nucleotides, making modification signatures highly context dependent ^36,37^. However, the ability of current algorithms to identify novel RNA modification sites *de novo* in biological samples remains largely untested, particularly given the diversity of modification chemistries, sequence contexts, and modification stoichiometries.

Recent comparisons of transcriptome wide modification maps derived using RNA-seq and DRS methodologies reveal substantial inconsistency between platforms ^29^. Rigorous benchmarking against orthogonal methods will be essential for interpreting computational modification calls in both RNA-seq and DRS data. One major barrier to evaluating both RNA-seq and DRS methodologies for modification mapping has been the requirement to compare predicted findings with established sites of modification in cells. While previously characterized rRNA and tRNA sites often serve this function ^38–41^, these controls are insufficient as they do not capture the diversity of sequence and structural contexts necessary to fully evaluate transcriptome-wide methodologies. Furthermore, modifications are incorporated within abundant rRNA and tRNA species at high stoichiometries, making it difficult to evaluate the quality of methods for detecting sub-stoichiometric sites in RNAs with >1000-fold less abundance (*e*.*g*. individual mRNAs). This challenge could be theoretically overcome by predicting modification sites in unmapped RNAs *de novo* using emerging sequencing technologies, and then examining each predicted site using the limited, but rigorous, field-standard approaches for RNA modification identification, such as LC-MS/MS and site-specific modification detection (e.g. SCARLET, CLAP, etc) ^13,14,42– 46^. While laborious, such an approach can provide a dependable ruler for assessing emerging transcriptome-wide methodologies. Here we combine Nanopore DRS, *in vitro* transcribed (IVT) RNA control, LC–MS/MS and CLAP to evaluate the RNA004 platform and Dorado modification calling using genomic RNA from the MS2 bacteriophage. MS2 is a well-studied positive sense, single strand RNA virus with a compact 3,569-nucleotide long genome^47^, providing a controlled and reproducible system for cross-validating modification calls and assessing potential false positives. We find that Dorado can identify candidate RNA modifications *de novo*, but that its predictions require careful filtering and orthogonal validation. Using such a pipeline, we discovered multiple sub-stoichiometric sites of pseudouridylation in the MS2

RNA genome. These results underscore the importance of integrating sequencing-based and biochemical approaches to improve the accuracy and interpretability of RNA modification detection. Additionally, they demonstrate that the rigorous application of orthogonal approaches can reveal new biological insights - even for molecules that have been well-studied for decades ^48^.

## RESULTS

### LC-MS/MS reveals a sparse MS2 modification landscape containing only pseudouridine

To evaluate the RNA modification landscape of the MS2 bacteriophage, we first used a nucleoside liquid chromatography-tandem mass spectrometry (LC-MS/MS) approach capable of quantifying 50 varieties of ribonucleosides in a single analysis ^10^. In this assay, purified MS2 RNA is enzymatically digested into its nucleoside building blocks, which are subsequently subject to analysis. The ratios of canonical nucleosides (A:G:U:C) that we measured closely matched the theoretical nucleotide composition of the MS2 genome **(Figure 1A)**. Together with parallel RNA-seq analysis of our samples (**Supplementary Figure S1**), these data confirm the high purity of the RNA preparation. Beyond the expected canonical nucleosides, LC-MS/MS detected a single abundant post-transcriptional RNA modification, pseudouridine (Ψ). Our measurements indicate that there is the equivalent of 1.1 ± 0.07 Ψ-modifications per full-length MS2 RNA molecule **(Figure 1B, Supplementary Table S1)**. We detected an additional eight types of modifications at trace levels consistent with the existence of minor RNA damage products or very low level tRNA contaminants (0.003 ± 0.001 to .037 ± 0.01 per MS2 molecule).

**Figure 1.**
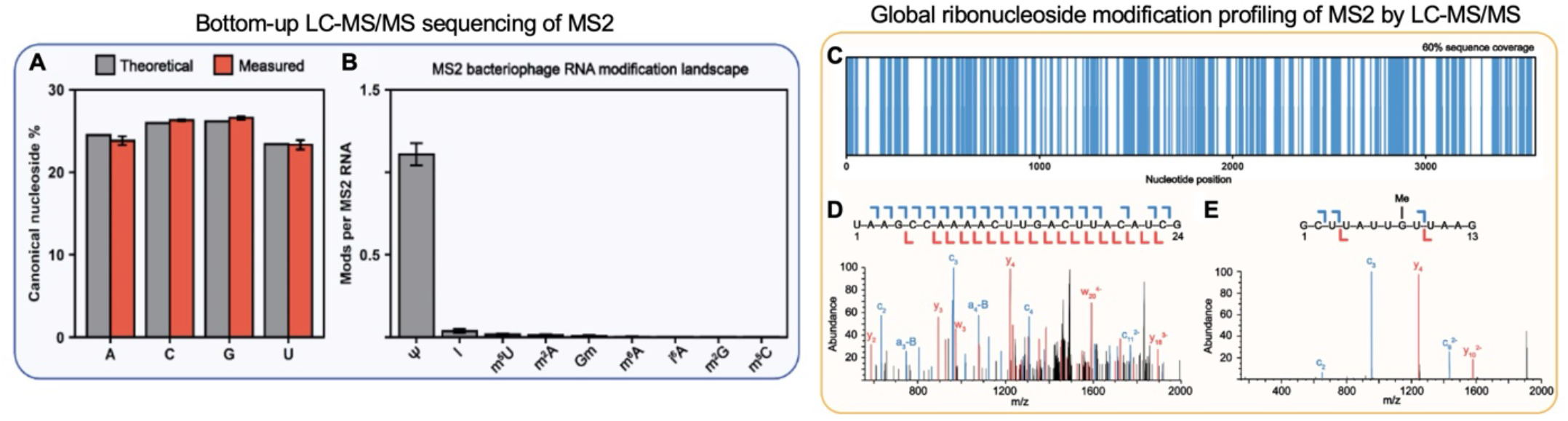
LC-MS/MS reveals Ψin MS2 RNA. **A**. The canonical nucleoside percentage determined by GRMP. **B**. The average number of modifications found per MS2 bacteriophage RNA for all modifications detected by GRMP. **C**. Sequence coverage map for the MS2 bacteriophage RNA using partial RNase T1 digestions, where blue bars represent a detected nucleotide position. **D**. Tandem mass spectrum of 24 nt oligonucleotide ion (1520.0092 m/z) with select fragment ions labeled. Blue bars represent fragment ions detected that contain the 5’ end of the selected oligonucleotide precursor. Red bars represent fragment ions detected that contain the 3’ end of the selected oligonucleotide precursor. **E**. Tandem mass spectrum of 13 nt oligonucleotide ion (1371.1813 m/z) that BioPharma Finder poorly identifies as methylated guanosine modification contain oligonucleotide fragment.

We next confirmed the sparsity of modifications in the MS2 genome by bottom-up LC-MS/MS RNA sequencing **(Figure 1C-E)**. Bottom-up LC-MS/MS is the current field-standard approach for detecting the sites of modifications within endogenous and synthetic RNAs ^14,46,49–52^. In this workflow, MS2 RNA was partially digested with RNase T1 to generate a complex mixture of oligonucleotides of lengths compatible with mass spectrometric analysis ^12,13,53^. The resulting oligonucleotides were separated by hydrophilic interaction liquid chromatography (HILIC) and analyzed by tandem mass spectrometry ^13^. Once identified, sequenced oligonucleotides were subsequently mapped back to specific regions of the MS2 genome ^13,53– 57^. In principle, LC–MS/MS sequencing can detect any RNA modification that produces a mass shift measurable by the mass spectrometer ^4,13^. Although analogous bottom-up workflows are widely used in proteomics, their application to RNA modification mapping remains challenging because RNA lacks broadly applicable digestion enzymes comparable to proteases used for protein sequencing ^58,59^. RNA LC–MS/MS workflows rely on base-specific nucleases, such as RNase T1, which cleaves after guanosine, or RNase A, which cleaves after pyrimidines. Because many RNAs contain frequent guanosines and pyrimidines, these enzymes often generate numerous short digestion products that cannot be uniquely mapped, resulting in low sequence coverage ^13,44,53,60^. This contrasts with trypsin and alternative proteases, such as Asp-N and Glu-C, used in bottom-up proteomics, that cleave at more restricted sets of sites and thereby generate peptides long enough to be uniquely assigned ^58,59^.

Using LC-MS/MS we obtained approximately 60% sequence coverage of the MS2 genome **(Figure 1C, Supplementary Table S2)** and confidently sequenced oligonucleotide RNase T1 digestion products up to 30 nt **(Figure 1 and Supplementary Figures S3, S4)**. During analysis, only a single modified position was identified by BioPharma Finder; however, further analysis revealed this modified position was a misidentified Na^+^ adduct (**Supplementary Figure S5)**. We did not obtain full sequence coverage of the MS2 bacteriophage RNA by bottom-up RNA sequencing **(Figure 1C)**, as expected. ∼60% sequence coverage is on par with a recent report, which also did not report any sites of modification in MS2^12^. Our sequencing findings support the nucleoside LC-MS/MS analyses we conducted in which Ψ was the only modification detected (**Figure 1A**). Unfortunately, one limitation of LC-MS/MS RNA sequencing is its inability to distinguish between modification structural isomers (e.g., ΨU, m^6^A/m^1^A), making it impossible to map Ψ sites using LC-MS/MS alone.

### Sequencing metrics for RNA002 and RNA004

Our characterization of the MS2 RNA genome by LC-MS/MS protocols revealed the presence of a modification (Ψ) that the technique cannot readily map. Nanopore DRS is commonly used to map Ψ across the transcriptome ^26,34,41,61^, so we sequenced MS2 RNA by nanopore DRS using both RNA002 and RNA004 platforms. In addition to providing further insight into the location of Ψ within the MS2 RNA modification profile, these experiments allowed us to benchmark current algorithms for *de novo* modification calling. The RNA002 sequencing was performed in 2019, yielded 536,618 reads with a read N50 of 3,455, while the RNA004 experiment, completed in 2024, yielded 22,937,016 reads with a read N50 of 3,589. After alignment to the MS2 reference genome, the RNA002 dataset contained 352,281 aligned reads (65.65% of total), whereas the RNA004 dataset had 17,354,168 aligned reads (75.66% of total). After filtering the RNA004 data for near full-coverage reads, a total of 8,895,271 reads were retained in the final BAM file.

To compare the quality of the data we collected with RNA002 and RNA004, we began by assessing alignment identity to the MS2 reference genome. RNA002, basecalled with Guppy, achieved a median identity of 89.61% for reads with at least 100 aligned bases. In contrast, RNA004, basecalled with Dorado v0.9.1, achieved a median alignment identity of 97.84 % using the same filtering criteria. These results demonstrate a notable improvement in alignment identity with the updated RNA004 chemistry (**Figure 2A**). To visualize base-level differences, we generated per-base substitution matrices. In RNA002, C-to-U and U-to-C substitutions were the most frequent, whereas G-to-C and C-to-G substitutions were the least common (**Supplementary Figure 6A**). A similar pattern was observed in RNA004, where U-to-C and C-to-U miscalls dominated, and G-to-C and C-to-G remained the least frequent (**Figure 2B)**. Compared to RNA002, the overall global miscall rate in RNA004 was reduced by at least 3%.

**Figure 2.**
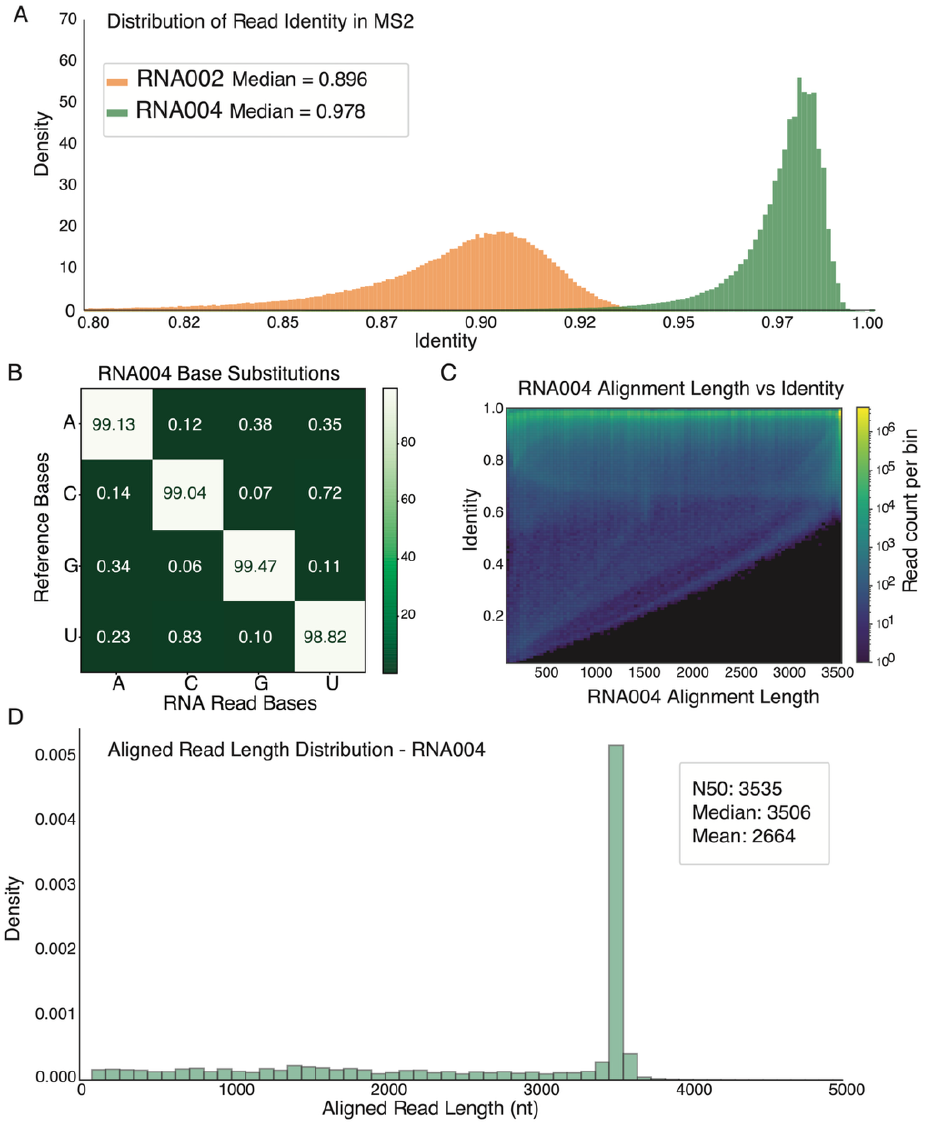
RNA004 direct RNA sequencing performance. **A**. Aligned read identity was calculated as a percentage of total matches over total matches, mismatches, insertions, and deletions in the aligned region for RNA002 and RNA004. **B**. RNA004 base-specific substitution matrix calculating the rate at which, given a reference base, each possible canonical nucleotide was observed in aligned regions of reads. **C**. 2D histogram of alignment length versus identity for RNA004. Aligned read length was calculated as the distance between the first and last aligned base of each read. **D**. Distribution of MS2 aligned read length in RNA004.

We computed the proportion of insertions and deletions (INDELs) in the aligned reads. RNA002 had a median INDEL rate of 0.0740, while RNA004 showed a lower median rate of 0.0141. The combination of reduced INDELs and an improvement in per-base substitution rates contributes to the higher identity observed for RNA004. We further evaluated the impact of read length on alignment identity in RNA002 and RNA004 using 2D histograms of alignment length versus identity (**Supplementary Figure S6B, Figure 2C**). RNA004 produced a median aligned read length of 3,506 and a mean of 2,664, showcasing its ability to cover nearly the full length of the MS2 genome (**Figure 2D**). Together, these results show that RNA004 Nanopore DRS, combined with updated basecalling, substantially improves MS2 read yield, alignment identity, full-genome coverage, and INDEL/substitution accuracy relative to RNA002, establishing a higher-quality dataset for downstream analysis.

### IVT-controlled nanopore DRS reveals limitations of U-to-C mismatch–based Ψ detection

We sought to apply two separate algorithms, ModID-p and NanoPsu, specifically designed to map Ψ. Both algorithms were trained on matched modified and unmodified datasets across NN(Ψ/U)NN 5-mers, and leverage signal differences that historically led to U-to-C mismatches in the RNA002 platform to identify putative Ψ sites ^34,61^. When applied to MS2 bacteriophage RNA NanoPsu predicted seven Ψ candidates with > 90% confidence, and ModID-predicted 64 sites **(Table 1)**. However, recent studies have cautioned against over-interpreting U-to-C mismatches as direct evidence of Ψ, especially with updated basecallers ^29^. We therefore repeated the analysis using an in vitro transcribed (IVT) MS2 RNA as a negative control. This IVT RNA shares the same sequence but lacks modifications and helps to reduce false positives by controlling for sequence context effects. With IVT filtering, ModID-p predicted 12 sites (U149, U771, U924, U1058, U1059, U1349, U2148, U2193, U2323, U2830, U3202, U3365) and NanoPsu retained four high-confidence sites (U924, U1059, U2521 and U3365), corresponding to U-to-C mismatches between the NCBI reference and our experimental consensus genome **(Table 1)**.

**Table 1.** Summary of *de novo* Ψ detection by orthogonal approaches. NanoPsu, Mod-p ID and Dorado algorithms were used to identify sites. The stoichiometry of putative sites was predicted by Dorado and measured by CLAP.

| Putative Ψ site | Algorithm |  |  | Stoichiometry measurement |  |
| --- | --- | --- | --- | --- | --- |
|  | NanoPsu | Mod-p ID | Dorado | CLAP | Raw Dorado Ψ |
|  |  |  |  | % modified | % modified |
| 149 | - | + | - | 0 | 0.81 |
| 771 | - | + | - | 0 | 0.8 |
| 924 | + | + | + | 23 ± 1 | 84.43 |
| 1058 | - | + | - | nd | 0 |
| 1059 | + | + | - | 12 ± 4 | 0 |
| 1349 | - | + | - | 0 | 0 |
| 2148 | - | + | - | nd | 0.14 |
| 2193 | - | + | - | nd | 0 |
| 2323 | - | + | - | 11 ± 8 | 0 |
| 2521 | + | - | - | 17 ± 6 | 0 |
| 2830 | - | + | - | 17 ± 0.1 | 0 |
| 3203 | - | + | - | 16 ± 4 | 0 |
| 3365 | + | + | - | 8 ± 3 | 0.1 |

Collectively, our LC–MS/MS and nanopore DRS analyses indicate that Ψ is incorporated into MS2 RNA. Despite these clear indicators that Ψ resides within MS2, given the diversity in the sites identified by DRS algorithms it remained unclear which sites were real. Furthermore, our nucleoside analysis indicated that 1.1-equivalents of Ψ are present per MS2 molecule, raising the question of what the levels of Ψ incorporation across the multiple sites predicted by those algorithms might be. To directly ask this question, we turned to CMC-RT and ligation assisted PCR analysis of Ψ modification (CLAP). CLAP is a quantitative RT– PCR-based method that measures Ψ stoichiometry in a site directed manner^42^. Ten candidate uridines were selected for investigation that were either predicted by a single algorithm (Mod-IDP: U149, U771, U1349, U2323, U2830 and U3202; NanoPsu: U2521), or by both algorithms (U924, U1059 and U3365) (**Table 1**). We validated Ψ incorporation at 70% of the tested sites: U924, U1059, U2323, U2521, U2830, U3202 and U3365 (**Table 1, Supplementary Figure S7**). All validated sites exhibited modest Ψ/U stoichiometries of ≤ 25%. Analysis of modified sequence motifs suggests the possibility that some sites may be incorporated by the pseudouridine synthases RluA or TruD (**Supplementary Figure S8**) ^62^. Notably, the summed Ψ stoichiometry across these sites was 1.04 ± 0.26 Ψ per MS2 genome, broadly consistent with the 1.1 ± 0.07 Ψ residues per MS2 molecule measured by nucleoside LC– MS/MS. These results reveal that bacteriophage MS2 is modified, raising the possibility that Ψ might have a biological consequence on phage lifecycle or evolution. Furthermore, our data suggest that while U-to-C mismatches patterns may be useful for identifying global trends, individual sites inferred from these signatures require orthogonal validation.

### Dorado predicts Ψ, m^6^A, m^5^C, 2′-OMeG, and Inosine in MS2 RNA

Dorado has emerged as the most widely used basecalling algorithm for identifying sites of RNA modification and currently predicts sites for several abundant modifications (m^6^A, Ψ Ψ, m^5^C, inosine, 2’-O-methylcytidine/adenosine/guanosine/uridine (Cm/Am/Gm/Um)) ^29^. To evaluate the consistency of modification detection across nanopore chemistries, we compared RNA004 Dorado calls to results from RNA002. To compare RNA004 Dorado Ψ calling with previously used methods in RNA002, the Modkit (an ONT modification analysis software) output was filtered to extract putative Ψ sites that had been identified by other approaches (**Table 1**). Of the 13 putative sites we detected in RNA002, one had modification occupancies greater than 10 % reported in the RNA004 Dorado results (Ψ924).

To broadly assess RNA modification detection with Dorado, we analyzed both global and site-level modification calls across the MS2 genome. In addition to Ψ, we used Dorado to call m^6^A, m^5^C, inosine, and 2′-Ome sites **(Figure 3A)**. Across all eight called modifications, Dorado identified 3,542 positions with valid full coverage. Among these, 10 sites showed a modification occupancy of 20% or higher **(Figure 3B, Supplementary Table S3)**. m^5^C accounted for five sites, followed by m^6^A (2 sites), inosine (1 site), 2′-OMeG (1 site), and Ψ (1 site). To enhance the specificity of Dorado modification calls, we applied an IVT-based correction strategy using a whole-genome IVT dataset^63^. IVT RNA, composed exclusively of canonical bases, provides a sequence-matched but unmodified control, making it an effective reference for identifying false positives. Using our pre-calculated 9-mer false positive rates, we refined the Dorado predictions by subtracting the corresponding IVT-derived occupancy from each biological call. Among these, m^5^C accounted for four sites, followed by m^6^A, Ψ, 2′-OMeG and inosine with 1 site each. Notably, about half of these sites fall near the 20 % occupancy threshold. Additionally, two sites (m^6^A926 and an m^5^C927) were located within 5 nucleotides of a high confidence Ψ (Ψ924), and another two sites (m^5^C3378 and m^5^C3379) were adjacent to each other.

**Figure 3.**
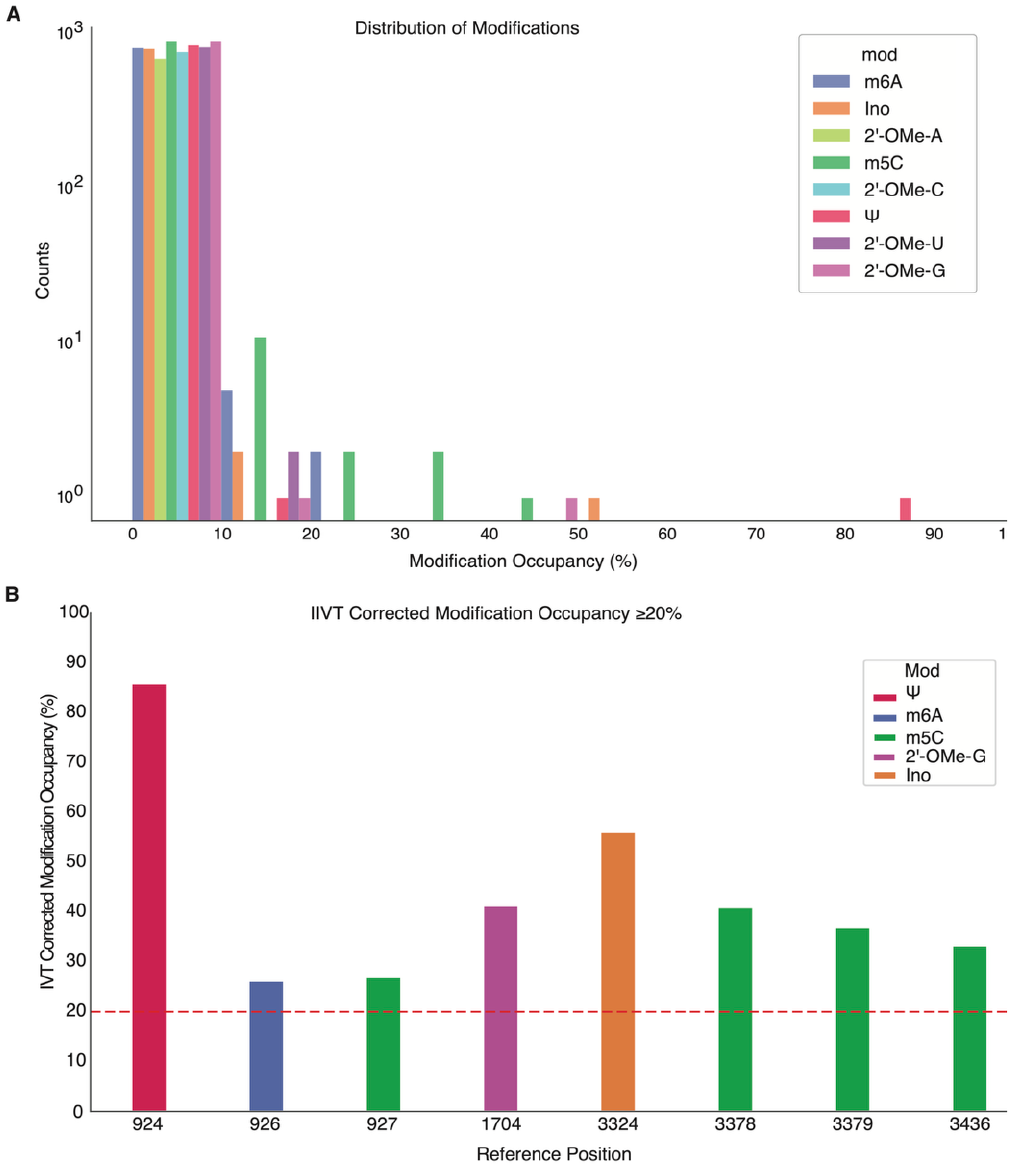
Dorado modification calling pattern in MS2. **A**. Distribution of putative modified sites for 8 types of modifications and their occupancy. Count of sites with valid full coverage reads. Counts per modification type were binned into 10% estimated modification occupancy windows starting at 20-30% occupancy and increasing by 10% increments. **B**. Dorado modified positions with >= 20% modification occupancy after subtracting IVT false positive rates (8 sites).

We next compared site-specific Ψ occupancy measured by CLAP with those predicted by Dorado. Dorado assigned high Ψ occupancy to U924, predicting an 84% modification fraction, and this position has also been predicted in a recent Nanopore analysis of the MS2 genome^64^. CLAP confirmed Ψ incorporation at the same position, but at substantially lower stoichiometry of 23%. Importantly, Dorado analysis detected four m^5^C, and one m^6^A, 2′-OMeG and inosine in the MS2 genome, which were all absent by nucleoside LC-MS/MS analysis (**Figure 1B**). Accordingly, we strongly recommend that sites of interest identified by Dorado be validated using orthogonal methods before biological interpretation (**Figure 4**).

**Figure 4.**
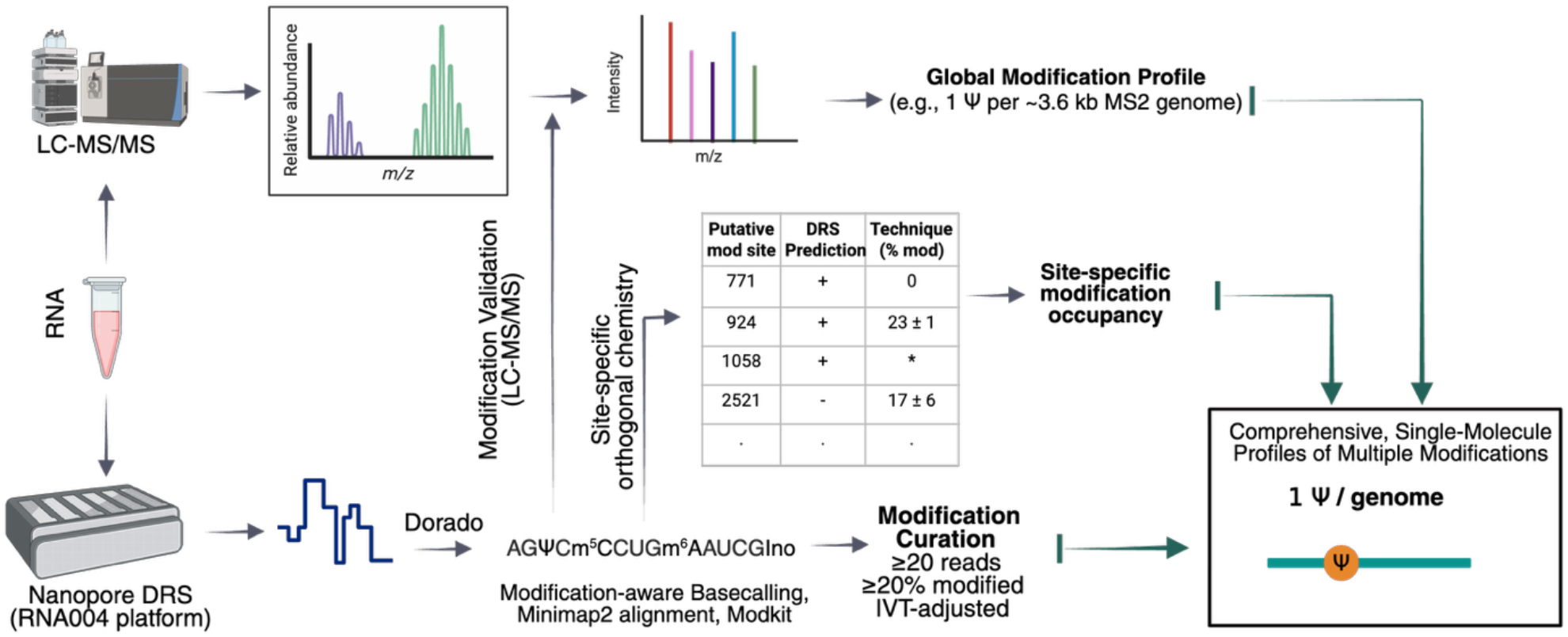
Guidelines for integrating orthogonal approaches to rigorously identify RNA modification sites *de novo*. We recommend that the modification profiles of samples be initially analyzed by both Nanopore DRS (or RNA-seq based methodologies) and LC-MS/MS nucleoside analysis. Putative mapped sites from DRS modifications should then be evaluated using nucleoside LC-MS/MS, and sites of interest must be further verified by orthogonal position-specific methods such as SCARLET, CLAP, primer extension, or LC-MS/MS sequencing.

## DISCUSSION

We combined LC–MS/MS with Nanopore direct RNA sequencing (DRS) and CLAP to characterize RNA modifications in the MS2 bacteriophage genome. We were able to identify multiple substoichiometric Ψ modification sites throughout the MS2 genome using DRS-guided analyses. However, orthogonal validation of individual sites by CLAP revealed that the stoichiometries inferred from Nanopore-based approaches were inaccurate, with substantial discrepancies between predicted and measured Ψ occupancies (**Table 1**). Our work is distinguished from available benchmarking studies that compare the findings of transcriptome-wide sequencing techniques to one another ^29,34,65^. Such studies, while essential to the field do not benchmark either *de novo* DRS or NGS approaches to an actual a “ground truth”, as we are able to do here through the dual application of LC-MS/MS and CLAP.

In general, we found that using the RNA004 sequencing chemistry and Dorado basecaller improved sequencing accuracy and throughput relative to RNA002 (**Figure 2**). This improved platform enables more informative analysis of modification-associated ionic current signals than previously available. Dorado identified candidate Ψ sites that were partially consistent with LC-MS/MS and CLAP measurements (**Table 1**). However, Dorado also produced high-confidence predictions for m^6^A, m^5^C, 2′-OMeG, and inosine at sites not supported by LC–MS/MS (**Figure 3**). Our observation of modification mis-calls by Dorado supported by a recent Nanopore DRS examination of MS2 RNA that predicted 8 sites of m^5^C modification ^64^. These results indicate that a subset of calls likely reflects model-dependent false positives or background signal misclassification. Collectively, our studies support the potential utility of RNA004-based DRS for *de novo* modification discovery, while highlighting the need for independent validation.

The level of precision required for modification mapping is higher than for sequencing canonical nucleotides because modifications are incorporated at low level (< 1% of bases transcriptome wide)^2^. As a result, the biological interpretation of a false positive modification call is more consequential than miscalling an abundant canonical nucleotide. Therefore, approaches that are > 99% accurate are required for reliable mapping of RNA modifications. This is illustrated by the fact that false positive rate for IVT RNA ranges between 0.0016-0.0086% for individual modifications, and Dorado still predicted seven aberrant modification sites in the 3,569-nucleotide MS2 genome ^29^.

Our data provide a rare benchmark dataset for assessing whether Nanopore algorithms can accurately identify RNA modifications and estimate their stoichiometry in native RNA. Many existing benchmarks rely on synthetic oligonucleotides, comparisons across sequencing platforms or perturbation of writer enzymes, each of which provides useful but incomplete ground truth. In contrast, we suggest that biologically isolated RNAs quantified by orthogonal approaches such as LC–MS/MS - and, where feasible, site-resolved methods such as CLAP or SCARLET - can more directly define the modification landscape against which Nanopore models are evaluated. Our results show that RNA004 chemistry improves the capacity to detect multiple modification classes at both per-read and per-site resolution, but also that modification calling can be sensitive to background error, model bias and the difficulty of quantitative occupancy estimation. Importantly, we find that neither Nanopore DRS or LC-MS/MS alone are able to fully characterize the MS2 RNA genome. However, the total Ψ content measured by nucleoside LC-MS/MS (∼1.1 Ψ/genome) was able be reconciled with multiple low-stoichiometry Ψ sites predicted by Nanopore DRS and validated by CLAP. Thus our findings provide not only a foundational dataset for validating sequencing approaches, but also suggest that viral RNAs may harbor a distributed landscape of low-occupancy modifications rather than a small number of constitutively modified sites.

Together, these data support Nanopore DRS as a powerful and rapidly advancing platform for native RNA modification analysis, while underscoring the need for rigorous validation. We find that broad trends observed across samples, sites or algorithms may be biologically informative, but individual modification calls, particularly quantitative estimates of occupancy, should be interpreted cautiously unless supported by orthogonal evidence. Incorporating IVT controls, benchmarking against LC– MS/MS and related methods, and prioritizing independent validation for sites of biological interest will improve the reliability of both RNA-seq and Nanopore-derived modification maps. As basecalling and modification-calling models continue to mature, this “trust but verify” framework will be essential for making quantitative RNA modification analysis robust, reproducible and broadly accessible for studies of post-transcriptional gene regulation (**Figure 4**).

## MATERIALS AND METHODS

### Expression and isolation of MS2 bacteriophage viral RNA

A 5 mL culture of NovaBlue(DE3) cells (Novagen) was grown in LB media supplemented with 10 mg/mL to maintain the F-plasmid. When cells reached the exponential phase (OD600 ∼0.4), the culture was inoculated with ∼1.5 × 10^10^ MS2 bacteriophages (MOI ∼10). The infected culture was incubated at 37°C for 16h with shaking (250 rpm). To isolate the phage RNA, the culture was centrifuged for 10 min at 1000 g to remove the cell debris and mixed with 5X PEG precipitation solution (2.5 M NaCl, 20% PEG w/v). Afte r 30 min incubation at +4°C, the supernatant was centrifuged for 15 min at 12’000 g to pellet the phage precipitate. The supernatant was discarded, and the phage pellet was resuspended in 140 μL of TE buffer. RNA was extracted from precipitated phage using QIAamp Viral RNA Mini Kit and eluted in 60 μL of AVE buffer (QIAGEN). Extracted RNA (10 uL) was reverse-transcribed with SuperScript IV First-Strand Synthesis System using the standard protocol (Invitrogen). A ten-fold diluted cDNA was used for RT-PCR with Q5 High-Fidelity DNA Polymerase (New England Biolabs) to generate overlapping amplicons that span the length of the phage genome. Sequences of oligonucleotide primers used for RT-PCR were as follows:

1F: GGGTGGGACCCCTTTCGG, 1R: TTTTTCTAGAGAGCCGTTGCCT;

2F: GGCCCAAATCTCAGCCATGC, 2R: CGTGTCTGATCCACGGC;

3F: GGCACAAGTTGCAGGATGCA, 3R: TGGGTGGTAACTAGCCAAGCAG.

### Global ribonucleoside modification profiling

To quantify up to 50 canonical and modified ribonucleosides, ribonucleoside LC-MS/MS was performed. The MS2 bacteriophage RNA (150 ng) was hydrolyzed to monoribonucleosides through a two-stage enzymatic digestion. First, the RNA was digested to ribonucleotide monophosphates using 300 U/μg Nuclease P1 (NEB 100,000 U/μL) overnight at 37 °C in 100 mM ammonium acetate (pH 5.5) and 100 μM ZnSO_4_. Following this, the nucleotides were dephosphorylated using 50 U/μg bacterial alkaline phosphatase (Invitrogen, 150 U/μL) for 5 hrs at 37 °C in 100 mM ammonium bicarbonate (pH 8.1) and 100 μM ZnSO_4_. Prior to each step, the enzymes were buffer-exchanged into their respective reaction buffers using a Micro Bio-Spin 6 size exclusion spin column (Bio-Rad). The resulting ribonucleosides were lyophilized and resuspended in 9 μL of water and 1 μL of 400 nM 15N_4_-inosine internal standard. The ribonucleosides were separated using a Waters Acquity HSS T3 column (1 × 100 mm, 1.8 μm, 100 Å) with a guard column at 100 μL/min at 35 °C on an Agilent 1290 Infinity II liquid chromatograph interfaced to an Agilent 6410 triple quadrupole mass spectrometer. Mobile phase A was water with 0.01% formic acid, and mobile phase B was acetonitrile with 0.01% formic acid. The MS conditions, LC gradient, and MS/MS transitions used were the same as previously published.

### Bottom-up LC-MS/MS of MS2 bacteriophage RNA

The MS2 bacteriophage RNA was digested using a mixture of RNase T1 (ThermoFisher) and quick calf intestinal phosphatase (Quick CIP, NEB). For fully digested MS2 bacteriophage RNA, 100 μg of RNA was digested with 10,000 U RNase T1 and 15 U Quick CIP for 30 min at 37°C in 100 mM ammonium acetate. For partial digested MS2 bacteriophage RNA, 250 μg of RNA was digested with either 500 U, 200 U, or 50 U RNase T1 and 30 U Quick CIP for 30 min at 37°C in 100 mM ammonium acetate. The digestion reactions were quenched by a phenol/chloroform extraction the addition of one volume of 125:24:1 phenol:chloroform:isoamyl alcohol mixture (Supelco, PN 77619). The resulting aqueous phase was washed once with chloroform prior to drying by lyophilization. Prior to use, the enzymes were buffer exchanged into 100 mM ammonium acetate prior to addition to reaction. Additionally, the 125:24:1 phenol:chloroform:isoamyl alcohol mixture was washed three times with an equal volume of water prior to use to remove excessive quantities of NaOAc that can affect downstream chromatography.

Prior to analysis, the RNA digestion products were resuspended in 10 μL of 200 mM ammonium acetate, and 5 μL was injected. The oligonucleotide digestion products were separated using a Waters Premier BEH Amide VanGuard FIT column (1 × 100 mm, 1.8 μm, 130 Å) at 250 μL/min at 55°C on an Agilent 1290 Infinity II Bio liquid chromatograph interfaced to a ThermoFisher Scientific Orbitrap Fusion Lumos mass spectrometer. Mobile phase A was 25 mM LC-MS grade ammonium acetate with 2.5 μM medronic acid (Agilent InfinityLab Deactivator Additive) in water (unadjusted pH), and mobile phase B was 25 mM LC-MS grade ammonium acetate with 2.5 μM medronic acid in 80:20 acetonitrile:water (unadjusted pH). The LC gradient is displayed in **Supplementary Table S4**. The separation was interfaced to the Orbitrap using a HESI-II source. The spray voltage was -2.8 kV, the sheath gas was 35, the aux gas was 10, the sweep gas was 0, RF lens was 50%, the ion transfer tube temperature was 350°C, and the vaporizer temperature was 350°C. Data acquisition was performed using data dependent acquisition (DDA) with the Orbitrap for both precursor and fragment ion scans at 60K and 30K resolution, respectively. MS1 scans were collected from 300-2000 m/z with a maximum injection time of 100 ms and normalized AGC target of 120%. The MS2 scans were collected from 150-2000 m/z with a maximum injection time of 200 ms and a normalized AGC target of 200%. Precursor ions with a 2 m/z isolation window were fragmented using collision induced dissociation (CID) with a collision energy of 35% and activation time of 10 ms. A single charge state between 2-6 with signal intensity above 25,000 was selected for each precursor for a dependent scan, and a maximum of five precursor ions were cycle. Following selection, the ion was excluded for 3 s.

### Analysis of bottom-up LC-MS/MS RNA sequencing of the MS2 bacteriophage RNA

LC-MS/MS data was analyzed using ThermoFisher BioPharma Finder 5.1 where oligonucleotide digestion products with greater than 90% confidence scores and maximum 10 ppm were designated a confident identification. All identified oligonucleotides were manually inspected for proper monoisotopic mass deconvolution and quality MS/MS spectra. Together, we detected 345 oligonucleotide digestion fragments resulting in approximately 60% sequence coverage of the MS2 bacteriophage RNA **(Figure 1)**, which is comparable to a previous attempt to sequence the MS2 bacteriophage using ion-pairing reverse phase chromatography that provides higher chromatographic resolution and MS sensitivity^12^. Using this technique, we could sequence confidently up to approximately a 30 nt digestion fragment **(Figure 1, S5, and Supplementary Table S2)** and detected only a single modified site following data processing and filtering (methylated adenosine). However, the detected modified site is of relatively low confidence for multiple reasons, despite being reported as 94% confidence by BioPharma Finder. Specifically, the methylated guanosine modification is detected on a 13 nt RNA; however, the retention time of the oligonucleotide is too early for an RNA of that length (20.5 min; coelutes with 4 nt digestion products). Further, the MS/MS spectra does not match well with the theoretical MS/MS spectra despite being relatively high abundance, unlike other confident oligonucleotide MS/MS spectra for RNAs of similar length. Following further analysis, the ion appears to be a Na+ adduct that was misidentified **(Supplementary Figure S5)**, from an unknown species likely arising from the sample preparation for a single sample.

### RNA002 direct RNA sequencing and modification analysis

MS2 bacteriophage RNA (4.6 μg) was incubated with 5 units of E. coli Poly(A) polymerase (NEB M0276S) at 37°C for 30 minutes to polyadenylate the 3’ end. RNA was purified using NGSpure magnetic beads. 600 ng of polyA MS2 RNA was used for sequencing. Library preparation of MS2 RNA included ligation of Nanopore adapters using T4 ligase and reverse transcription of a cDNA scaffold using Superscript III reverse transcriptase. Direct RNA sequencing was performed using a MinION (Oxford Nanopore Technologies) and the direct RNA sequencing kit (SQK-RNA002) per manufacturer’s instructions.

IVT RNA was generated by reverse transcription of Native RNA into overlapping segments using 3 MS2 primer sets and Superscript III RT. DNA fragments were transcribed using the T7 high yield transcription kit to generate IVT MS2 RNA for nanopore sequencing. IVT RNA was sequenced as described above using SQK-RNA002 kit per manufacturer’s instructions. Native and IVT MS2 RNA reads were used to predict base modifications using NanoPsu version 3.8.8 to detect Ψ. For ΨΨsites were detected in RNA002 data using Mod-p ID v1.0.0. The U-to-C mismatch frequency was calculated by comparing native RNA sequences against IVT controls.

The significance (p-value) of each U site was determined based on the number of reads and the mismatch level.

### Ligation-assisted PCR analysis of Ψ modification (CLAP)

CLAP was conducted as previously described^42^. Primer sequences were designed using Primer3^66^ and checked for specificity using primer BLAST. The CLAP primers (designed by primer3 for U149, U771, U924, U1059, U1349, U2323, U2521 and U2830), adaptor, and splint sequences are listed in **Supplementary Table S5**.

### RNA004 direct RNA sequencing, basecalling, alignment

We used 600 ng of MS2 RNA input following the DRS library preparation protocol with SQK-RNA004 kit<u>^18^</u>. The library was sequenced using an RNA004 flow cell on a PromethION platform for 72 hours. The resulting Pod5 files were basecalled using Dorado basecaller v1.0 with the super accuracy model (rna004_130bps_sup@v5.1.0) and modification calling enabled for eight modifications simultaneously.

~~~
dorado basecaller sup --modified-bases
pseU_2OmeU 2OmeG m5C_2OmeC
inosine_m6A_2OmeA
~~~

The resulting basecalled BAM file was converted to FASTQ while preserving modification tags.

~~~
samtools fastq -T’*’
~~~

The FASTQ file was then aligned to the NCBI phage MS2 reference genome (NC_001417.2) using minimap2, with - y to retain modification tags.

~~~
minimap2 -uf -k 14 -ax splice -y
~~~

The resulting BAM file was filtered for primary alignments, sorted, and indexed using Samtools.

~~~
samtools sort -o | samtools index
~~~

### Preprocessing and statistical analysis

To extract base-level statistics against reference genome positions, we first ran Pysamstats on the aligned and sorted BAM file using the following command:

~~~
pysamstats --type=variation_strand --
fields=“chrom,pos,ref,deletions,deleti
ons_fwd,deletons_rev,insertions,inser
tions_fwd,insertions_rev,A,A_fwd,A_rev
,C,C_fwd,C_rev,T,T_fwd,T_rev,G,G_fwd,G
_rev”
~~~

Per-read identity was calculated as the number of matches divided by the sum of matches, mismatches, insertions, and deletions for each read aligned to the reference. To ensure reliable measurements, we restricted the identity calculation to reads with an aligned length of at least 100 bases, defined as the difference between the reference end and start positions.

To calculate base substitutions, we processed the sequencing data by reading the Pysamstats output. At each genomic position, we extracted the reference base and the observed counts for each base (A, C, G, T). Positions with a total base count of at least 20 reads were considered valid for analysis. For each valid site, we compared the reference base to the most frequently observed base, recording how often one base was substituted for another. Additionally, we calculated the overall substitution fractions by summing the observed base counts for each reference nucleotide and computing the relative frequency of each substitution type.

### Dorado modification calling

For analyzing modification calls, the BAM file was further filtered using Pysam to retain only reads with near full coverage of the MS2 genome (excluding the last 20 bases) and a mapping quality score of 60. To identify putative modification sites in the RNA004 dataset, the resulting sorted and filtered BAM file was processed using ONT’s Modkit tool17, which extracts predicted modification information based on basecalling signals. We used modkit sample-probs to get the filter threshold based on the 10th percentiles of data.

~~~
modkit sample-probs --sampling-frac
0.1 --hist --seed 42
modkit pileup --sampling-frac 0.1 --
filter-threshold T:0.94 --filter-
threshold C:0.85 --filter-threshold
A:0.77 --filter-threshold G:0.95 --max-
depth 100000 --seed 42
~~~

The resulting modification calls were mapped back to the reference genome to exclude sites likely arising from mismatch errors. Further filtering was applied to retain only those positions with at least 20 reads of coverage and a modification confidence of 20% or higher, helping to reduce false positives.

~~~
modkit extract calls --mapped-only --
ref
~~~

### Subtracting false positive rates and re-filtering with Whole Genome IVT Data

Using a 9-mer false positive rate table from the whole genome IVT dataset, we calculated the reported modification occupancy of each site in the Modkit pileup. After subtracting the false positive rate, we refiltered for sites that continued to have a Dorado modification occupancy over 20 % and a minimum of 20 valid full read coverage.

~~~
python IVT_fp_correction.py \
 --modkit input_bedMethyl.tsv \
 --reference reference_genome.fa \
 --errortable
./error_tables/GM12878_genomic_IVT_ref
match_9mer_all_threshold.dorado_1.0.ts
v.gz
 --outpath corrected_bedMethyl.tsv
 --valid_only \
 --filter_mismatch \
 --filter_kmer
~~~

## ACKNOWLEDGEMENTS

We thank Maddy Zamecnik and Dr. Brandon Ruotolo for their careful reading of the manuscript and suggestions. We also the following funding sources for their support: National Science Foundation (CAREER 2045562 to K.S.K, Graduate Research Fellowship Program to J.D.J.), National Institute of Health (R01 HG013876 to K.S.K. and M.J.). Research reported in this publication was supported by the Office of the Director, National Institutes of Health under Award Number S10OD021619.

## COMPETING INTERESTS

The authors declare no competing interests.

## Notes

### Competing Interest Statement

The authors have declared no competing interest.

