## Supplementary Figures for "Integrating mass spectrometry with Nanopore direct RNA sequencing for *de novo* modification profiling of bacteriophage MS2"

**SUPPLEMENTARY FIGURE 1**

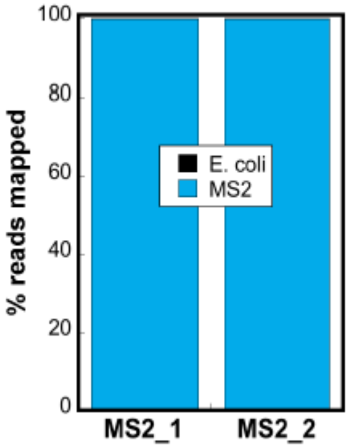


**Supplementary Figure S1: MS2 bacteriophage purity analysis by RNA-seq.** For two replicates, the distribution of reads corresponding to either the MS2 bacteriophage (blue) or *E. coli* RNA (black) are displayed.

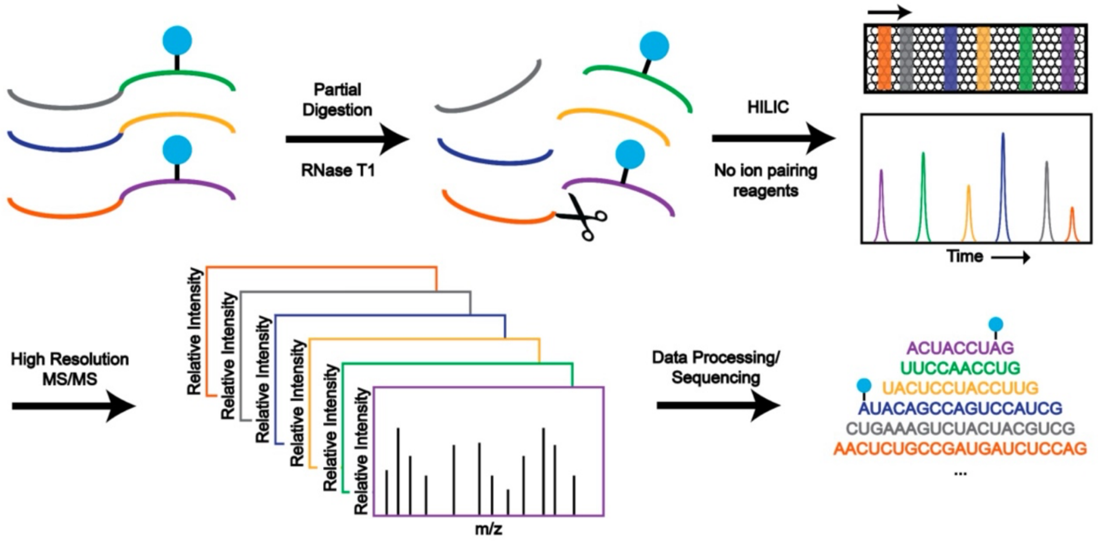


**Supplementary Figure S2: Supplemental Figure S3: Bottom-up LC-MS/MS RNA sequencing workflow.** The MS2 bacteriophage RNA was partially digested with RNase T1 under four separate digestion conditions. The resulting oligonucleotide digestion fragments were separated using HILIC and detected by high resolution MS/MS on an Orbitrap Fusion Lumos. The resulting collision induced dissociation (CID) fragmentation spectra were sequenced using BioPharma Finder 5.1 to identify modified positions.

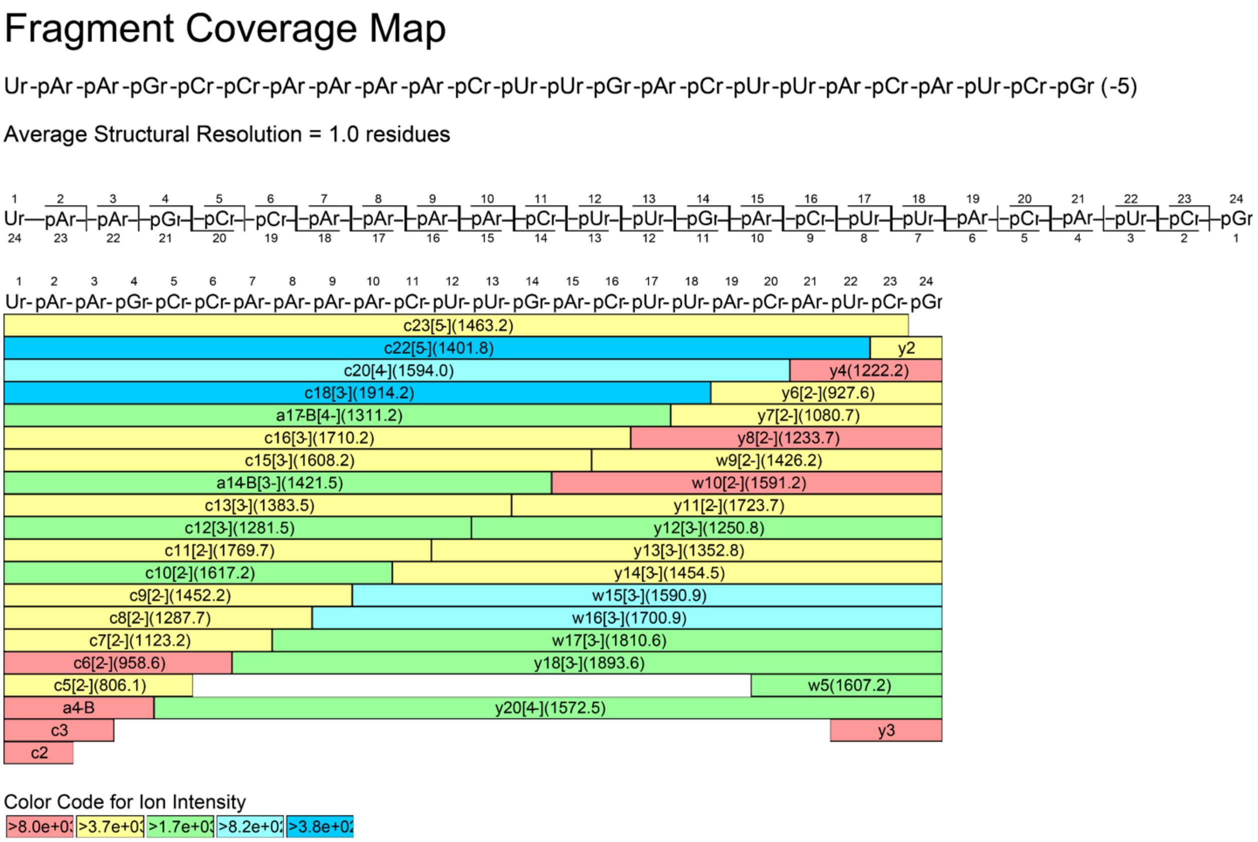


**Supplementary Figure S3: Fragment coverage map of 24 nt RNA detected by tandem mass spectrometry.** Identified oligonucleotide fragment ions in the CID MS/MS spectrum are shown. Each bar is labeled with the type of fragment ion, charge state of the fragment ion, and the m/z of the fragment ion in the MS/MS spectrum.

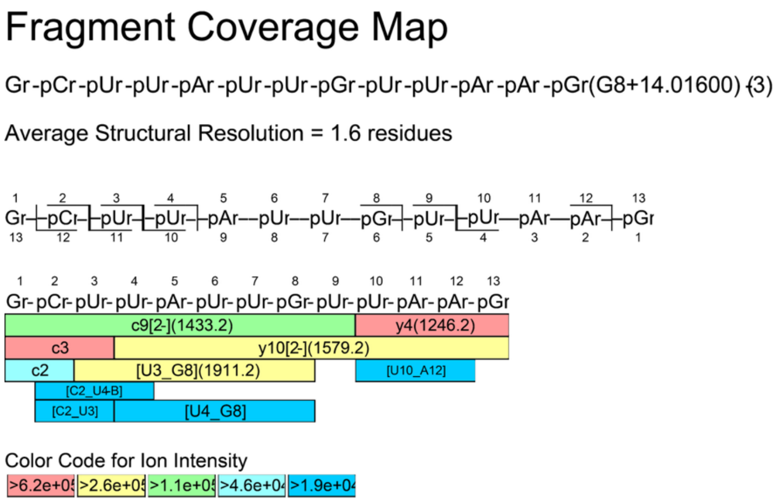


**Supplementary Figure S4: Fragment coverage map of 13 nt oligonucleotide digestion product that was identified to contain methylated guanosine modification.** Identified oligonucleotide fragment ions in the CID MS/MS spectrum are shown. Each bar is labeled with the type of fragment ion, charge state of the fragment ion, and the m/z of the fragment ion in the MS/MS spectrum. BioPharma Finder specifies that methylated guanosine modification is likely present on nucleotide position 8.

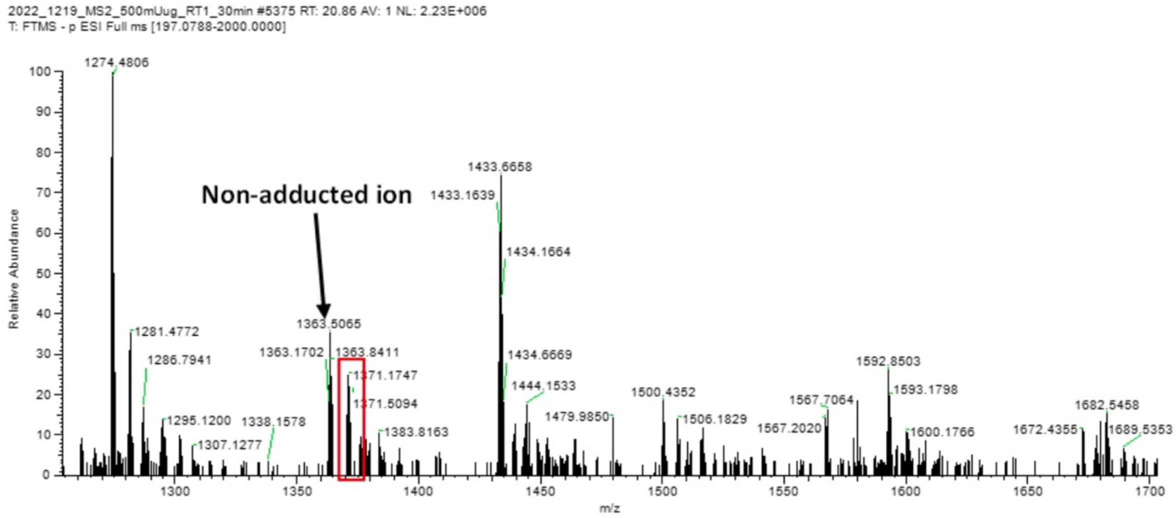


**Supplementary Figure S5: MS/MS sequencing analysis with BioPharma Finder that identified methylated modification site.** Tandem mass spectrum of 13 nt oligonucleotide ion (1371.1813 m/z) is displayed (bottom panel) in comparison to a theoretical MS/MS spectrum based on the RNA sequence (top panel). BioPharma Finder compares the two spectra to calculate the confidence score of the identification, and BioPharma Finder reports a 94% confidence score that this oligonucleotide contains a methylated guanosine modification. Blue bars represent fragment ions detected that contain the 5' end of the selected oligonucleotide precursor. Red bars represent fragment ions detected that contain the 3' end of the selected oligonucleotide precursor.

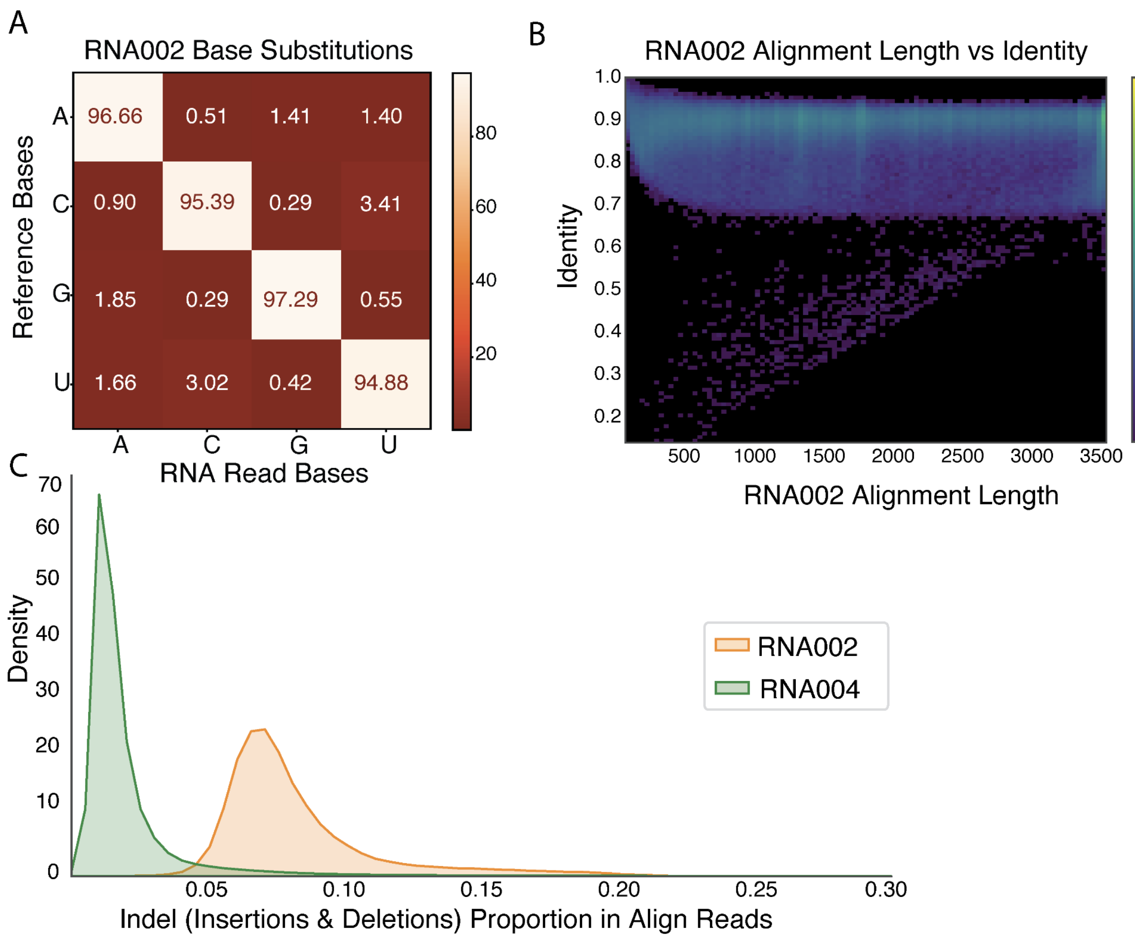


**Supplementary Figure S6. RNA002 direct RNA sequencing performance. (A)** RNA002 base-specific substitution matrix calculating the rate at which, given a reference base, each possible canonical nucleotide was observed in aligned regions of reads. **(B)** 2D histogram of alignment length versus identity for RNA004. Aligned read length was calculated as the distance between the first and last aligned base of each read. **(C)** The proportion of insertions and deletions in RNA002 and RNA004.

A.

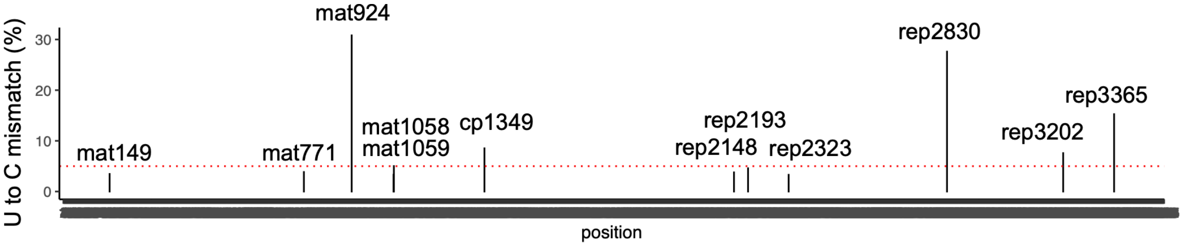


B.

1. mat149 CCUUG

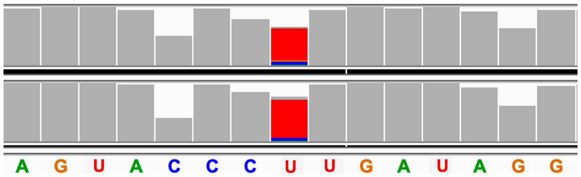


2. mat771 UAUGA

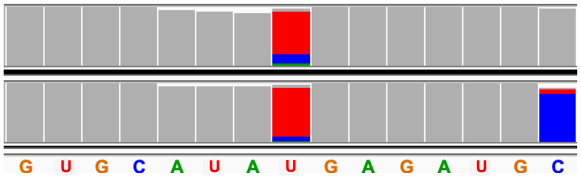


3. **mat924 UUUUA**

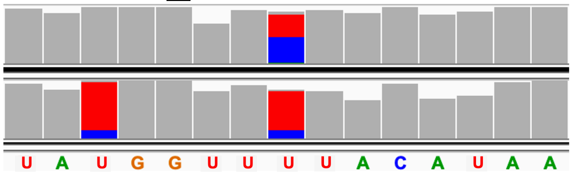


4. mat1058 CCUUA

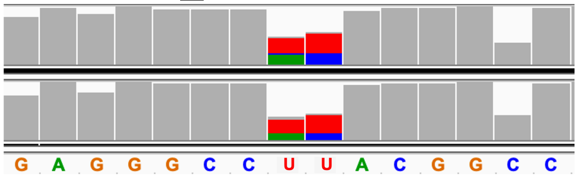


5. mat1059 CUUAC

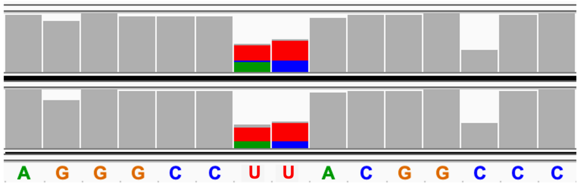


6. cp1349 UUUAC

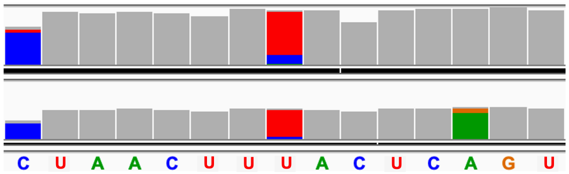


7. rep2148 UCUCC

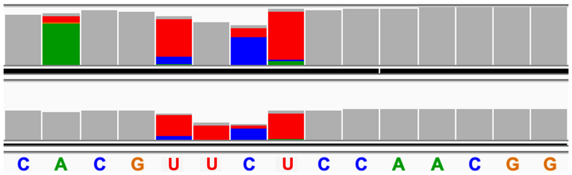


8. rep2193 CUUAC

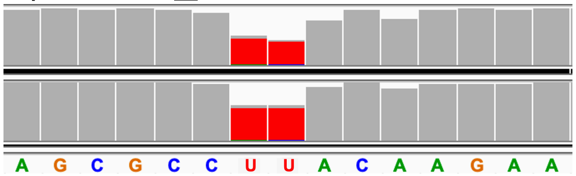


9. rep2323 CGUUG

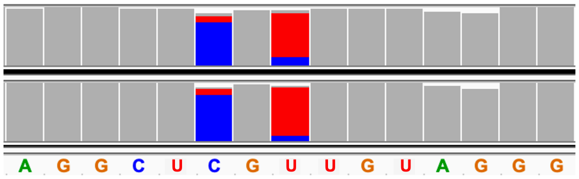


10. rep2830 ACUUG

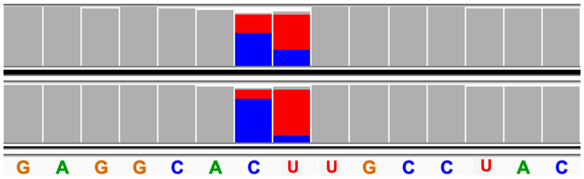


11. rep3202 UCUUG

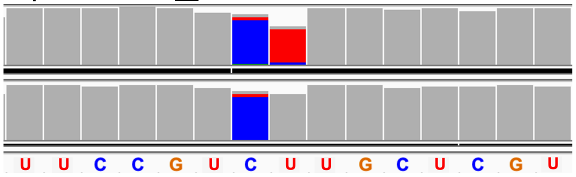


12. rep3365 CCUCA

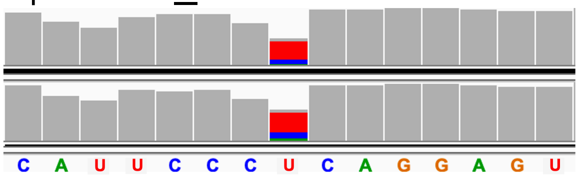


**Supplementary Figure S6: Predicted Ψ Sites Using PsiNanopore from ModID-p.** A. Predicted Ψ positions in MS2 RNA and individual U-to-C mismatch ratios (rep3202, rep3365 p-value < 0.01). B. Snapshots from the Integrated Genome Viewer (IGV) of aligned nanopore reads to the MS2 genome at predicted Ψ sites. Correctly aligned bases are shown in gray, while miscalled bases are represented in colors (cytidine, blue; adenine, green; guanine, orange; uridine, red) (the above one is native RNA, the below one is IVT). The genomic reference sequence is converted to the sense strand and displayed as RNA for clarity.

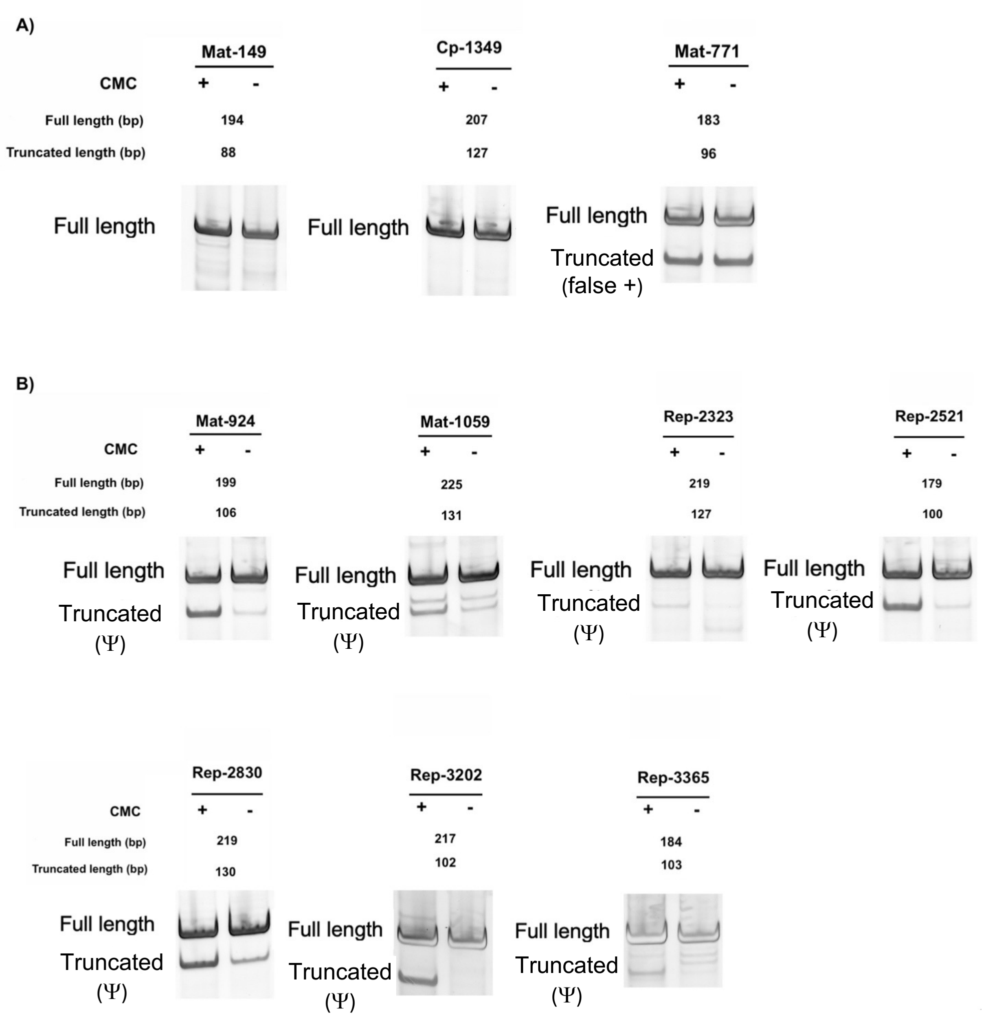


**Supplementary Figure S7. CLAP assays results examining 10 predicted sites of Ψ-incorporation in MS2. (A)** No Ψ incorporation was confirmed at predicted positions 149, 1771, and 349. **(B)** Ψ levels were measured at positions 924, 1059, 2323, 2521, 2830, 3202, 3365. Stoichiometries were calculated using the difference of % of total signal in the truncated band in CMC(-) and CMC(+) conditions.

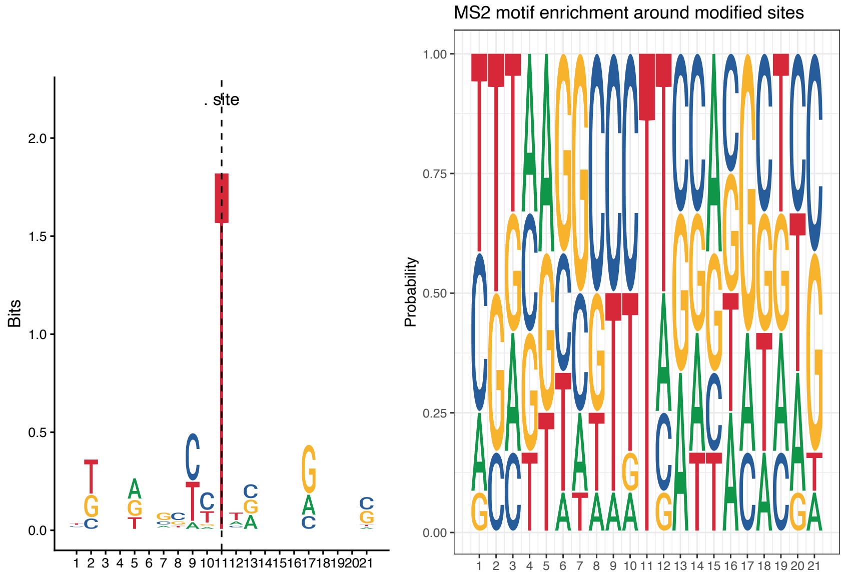


**Supplementary Figure S8. Sequence motif enrichment around modified Ψ (position 11).** Known sequence motifs for pseudouridine synthases include: GUΨCNANNC (TruB, bacterial); ΨUACA (RluA, bacterial); UNΨAR (Pus7 in eukaryotes)
